# Mono and Multidomain Anti-phage Defense Toxins of the RelE/ParE Superfamily

**DOI:** 10.1101/2025.01.28.635273

**Authors:** Kenn Gerdes

**Affiliations:** Independent Researcher at Voldmestergade, Copenhagen, Denmark

**Keywords:** RelE, ParE, Toxin, mRNase, Antitoxin, Anti-phage, Defense, Ribosome Dependency

## Abstract

Toxin-antitoxin (TA) modules are widely distributed across prokaryotes, often existing in large numbers despite their associated fitness costs. Most type II TA modules are bicistronic operons encoding a monodomain toxin and its cognate protein antitoxin. The RelE/ParE superfamily encompasses toxins with a conserved Barnase-EndoU-ColicinE5/D-RelE (BECR) fold, yet their cellular targets differ remarkably: RelE toxins function as ribosome-dependent RNases, while ParE toxins act as DNA gyrase inhibitors. Using a comprehensive bioinformatics approach, this study analyzed 13 BECR-fold toxin families as classified in the Pfam database. Intriguingly, the ParE family was found to include a subcluster of mRNA-cleaving toxins, challenging its conventional role as solely DNA-targeting. This study identified a novel tripartite operon encoding a PtuA-like defense ATPase, a homolog of type IV restriction endonucleases, and a RelE homolog, suggesting a coordinated role in defense mechanisms. Multidomain BECR-fold toxins, including transmembrane variants, were also discovered, extending the functional repertoire of type II TA modules to membrane-associated systems. These findings clarify the evolutionary relationships and functional diversity within the RelE/ParE superfamily, and discovers novel, putative defence systems that can now be investigated experimentally.

**Importance:** Toxin - antitoxin modules play critical roles in prokaryotic survival and adaptation, contributing to genome stabilization and defense against phages and invading plasmids. The RelE/ParE superfamily exemplifies the structural and functional diversity of these systems, with members targeting distinct cellular processes, such as translation and DNA supercoiling. By elucidating the relationships among the 13 BECR-fold toxin families, this study enhances our understanding of microbial resistance mechanisms and reveals potential new opportunities for research into prokaryotic defense and regulation. These insights may have significant implications for medical and biotechnological applications, particularly in understanding bacterial responses to genetic invaders.

## Introduction

Toxin – antitoxins (TA) modules are nearly ubiquitous across the prokaryotic world, encoded by plasmids, phages, integrons, defence islands and elsewhere on chromosomes. Initially identified for their role in plasmid stabilization through post-segregational killing of plasmid-free cells (1, 2), subsequent studies have revealed their widespread chromosomal presence, often in surprisingly high numbers (3–5). This suggests additional roles beyond plasmid maintenance. For instance, transcription of some TA loci is induced by nutritional stresses and one locus in particular was indicated to be involved in the stringent response via interaction with the ribosome, linking TA systems to bacterial physiology and survival (6, 7). Furthermore, TA loci have been associated with bacterial persistence, antibiotic tolerance, pathogenesis and biofilm formation, though these connections may depend on the experimental methodologies used (8, 9). Recent evidence indicates that TA modules frequently function as prokaryotic defense mechanisms that limit the invasion of phages, plasmids, and other foreign genetic elements (8, 10–26).

Toxin – antitoxin modules typically encode two components: a toxin that inhibits cell growth or induce cell death and an antitoxin that neutralizes the toxin’s activity. TAs are classified into different types based on how the antitoxin neutralizes the toxin. Type I and III modules use antisense RNAs to block toxin mRNA translation or directly bind and neutralize the toxin, respectively. Type II TAs, the most common type, encode protein antitoxins that directly bind and neutralize cognate toxins. These systems are further classified into families based on structural and sequence similarities among toxins. Typically, but not universally, toxins within a family share similar molecular mechanisms. A recurring theme involves growth inhibition through translation interference by mRNA, tRNA, or rRNA cleavage, tRNA modification, or translation factor inactivation (27–34).

Among translation-targeting toxins, several families, known as mRNA interferases or mRNases, inhibit translation by cleaving mRNA. For example, the HicA toxin family consists of small ribonucleases with double-stranded RNA-binding domains (35), while the RelE superfamily of RNases exhibit the <u>B</u>arnase-<u>E</u>ndoU-<u>C</u>olicinE5/D-<u>R</u>elE like nuclease (BECR) fold, comprising an N-terminal α-helical segment followed by a domain of three or four antiparallel β-sheets (36, 37). The model toxin RelE of *E. coli* K-12, a mono-domain RNase, adopts a β-α-α-β-β-β-α topology with a compact β-sheet core flanked by three α-helices (**Figure 1A, 1B**). Free RelE is inactive but binds to the ribosomal A-site loaded with mRNA codons (**Figure 1C**), where it cleaves the mRNA between the second and third codon nucleotides (38–41). RelE homologs, including multidomain versions, also cleave A-site codons along the mRNA between the 2^nd^ and 3^rd^ codon nucleotides or, in rare cases, between individual codons (42–49). RelE of *E. coli* K-12 can also cleave A-site codons of archaeal and eukaryotic ribosomes (39, 50).

**Figure 1.**
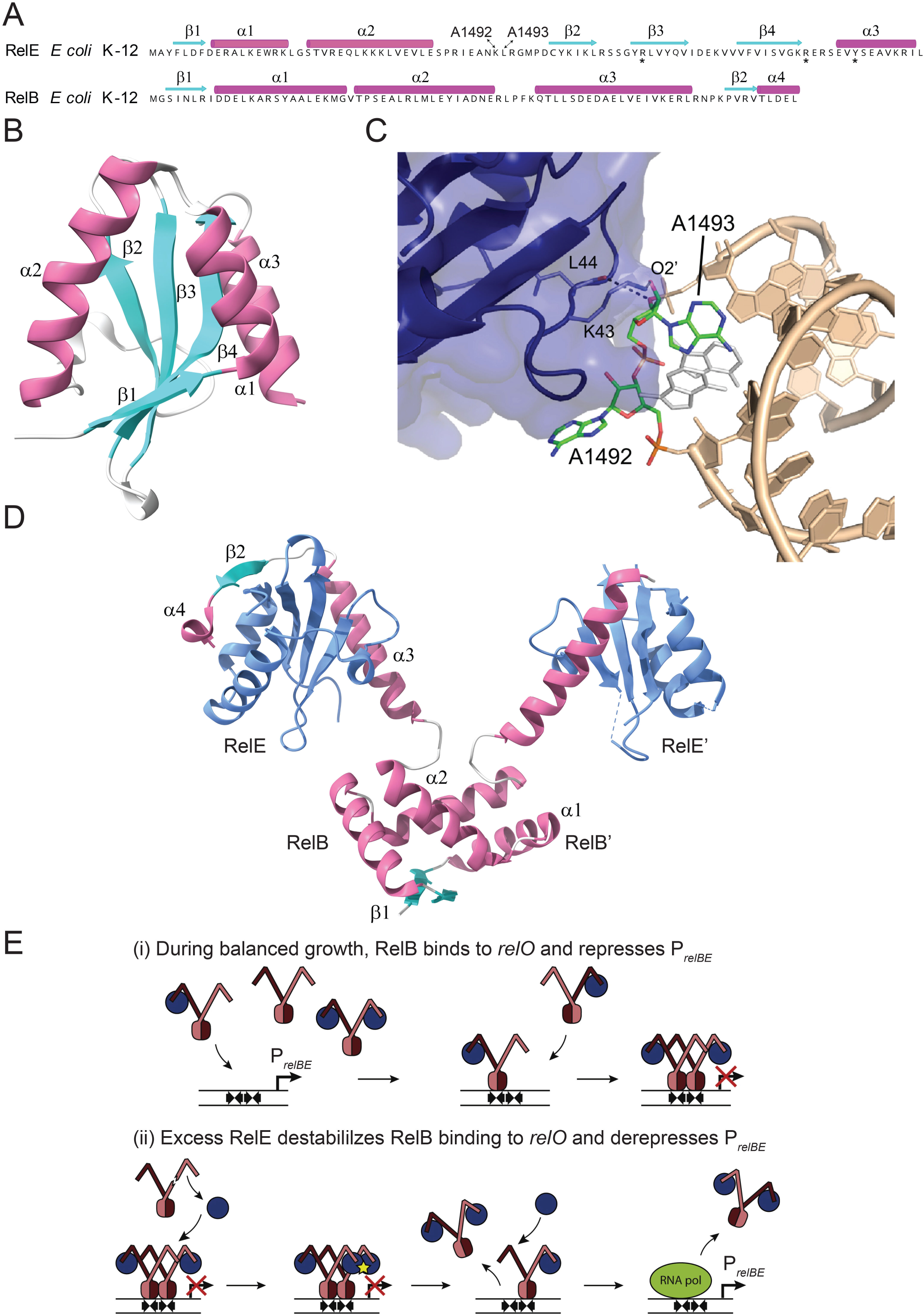
Components and regulatory features of the *relBE* module of *E. coli* K-12 A. Primary and secondary structural features of RelE (CAA26251.1) and RelB (CAA26250.1). The amino acid sequences are shown with secondary structure annotations: β-sheets are represented in cyan, and α-helices are in magenta. Adapted from (51). B. Tertiary structure of free RelE, featuring its compact conformation. The β-sheets (cyan) form a stable core flanked by α-helices (magenta) (51, 112, 113). C. Structural interactions between RelE (blue, represented with a semi-transparent surface and critical residues shown as sticks) and the decoding center of the ribosomal A-site. Bases A1492 and A1493 in helix 44 of 16S rRNA (wheat, sticks) undergo conformational changes upon RelE binding. Key interactions involve lysine 43 (K43) and leucine 44 (L44) from the loop between α2 and β2 of RelE, which contact the decoding center adenosines (green sticks). These interactions disrupt normal decoding and facilitate mRNA cleavage at the A- site (113). The unbound conformation of the decoding center adenosines is shown in white for comparison. D. Tertiary structure of the RelB₂E₂ quaternary complex. The two RelE monomers (blue) are neutralized by the RelB antitoxin dimer, composed of β-sheets (cyan) and α-helices (magenta) (51). E. Regulatory model of *relBE* operon transcription by conditional cooperativity. Upper panel: During balanced cell growth, RelB is in excess, forming RelB₂ dimers (brown and dark brown). Two RelB₂E heterotrimers bind cooperatively to adjacent *relO* operator sites, forming a RelB₂E-RelB₂E heterohexamer that represses *relBE* operon transcription. Lower panel: When the RelE:RelB ratio shifts in favor of free RelE (e.g., during stress or phage infection), free RelE interacts with the unoccupied C-terminal tails of RelB within the heterohexamer, inducing steric clashes. This destabilization results in the release of the RelB₂E₂ complex from the *relO* operators. Additional free RelE may bind and displace remaining complexes, enabling transcriptional derepression (51, 53). An alternative model for *relBE* regulation has been proposed by (114).

The cognate antitoxin RelB prevents RelE activity by binding directly to the toxin and blocking ribosomal A-site interactions. RelB dimerizes via its N-terminal Ribbon-Helix-Helix (RHH) DNA- binding domain, and each C-terminal segment wraps around a RelE monomer (**Figure 1D**) (51).

RelB dimers bind cooperatively to the *relBE* operator, repressing operon transcription (51–53).

Operon regulation follows conditional cooperativity: a high RelB:RelE ratio represses transcription, while excess RelE destabilizes RelB binding, derepressing transcription (**Figure 1E)** (51).

TA modules encoding BECR-fold toxins belong to the RelE/ParE superfamily (54). While ParE toxins, which inhibit DNA gyrase, share the BECR fold, they lack essential catalytic residues typical of BECR-fold RNases. At least 13 distinct BECR-fold toxin families have been identified, yet inconsistent naming conventions complicate their classification. This work employs bioinformatics and database analyses to provide a comprehensive overview of the 13 monodomain BECR-fold toxin families and novel multidomain variants. The findings strongly support the role of both mono- and multidomain BECR-fold toxins in anti-phage defense, advancing our understanding of their functional diversity and evolutionary significance.

## Results and Discussion

### BECR-Fold toxin families encoded by type II toxin-antitoxin modules

Extensive research into Type II toxin-antitoxin (TA) systems has uncovered a vast array of modules distributed across prokaryotic organisms (18, 55, 56). *Mycobacterium tuberculosis* and *Microcystis pseudomonas* strains harbor up to 100 and 130 Type II TA modules, respectively, underscoring their prevalence and potential biological significance (25). The model organism *E. coli* K-12 is equipped with at least 12 different Type II TA modules, seven of which encode monodomain BECR-fold toxins (40, 49, 57–60). These seven toxins, identified by genetic analyses, protein sequence similarity searches, or serendipitous discovery, include those with names such as RelE, HigB, MqsR, YafQ, YafO, YoeB, and YhaV.

The Pfam database classifies 13 distinct BECR-fold monodomain toxin families within typical Type II TA modules (**Table 1**). Among these, 11 families have been experimentally confirmed as RNases that cleave mRNA, similar to the well-characterized RelE of *E. coli* K-12. In contrast, the ParE family, despite sharing the BECR fold with RelE-like toxins, lacks RNase activity and instead inhibits DNA gyrase (54, 61, 62).

**Table 1.** The RelE/ParE Superfamily of BECR-fold Toxins Encoded by Type II Toxin – Antitoxins.

| Toxin Family name in Pfam | Pfam ID<sup>a)</sup> | Number of entries in Pfam<sup>b)</sup> | ID used here | Toxin names in Pfam<sup>c)</sup> | Cellular Target of Toxin | Ribosome Dependency | GO<sup>d)</sup> | References |
| --- | --- | --- | --- | --- | --- | --- | --- | --- |
| RelE | PF06296 | 6K | PF_RelE | HigB2, RelE, RelE/ParE, StbE, | mRNA degradation | Dependent | T/A | (47) |
| YoeB | PF06769 | 12K | PF_YoeB | YoeB, Txe, RelK, | mRNA degradation | Dependent | A/T | (40, 45, 68, 70, 115) |
| HigB | PF09907 | 8K | PF_HigB | HigB, HigB-like, RelE, YgjN, MenG, RelE/ParE | mRNA degradation | Dependent | T/A | (57, 66) |
| HigB-like | PF05015 | 14K | PF_HigB-like | HigB-like, HigB, HigB-1, RelE/ParE, RelE, StbE | mRNA degradation | Dependent | T/A | (43, 47, 71) |
| MqsR | PF15723 | 1k | PF_MqsR | MqsR, YgiU, | mRNA degradation | Independent<sup>e)</sup> | T/A | (57) |
| YafQ | PF15738 | 9K | PF_YafQ | YafQ, RelE, StbE, HigB, | mRNA degradation | Dependent | A/T | (44, 60, 116) |
| YafO | PF13957 | 1K | PF_YafO | YafO | mRNA degradation | Dependent<sup>f)</sup> | A/T | (57) |
| YhaV | PF11663 | 2K | PF_YhaV | YhaV, RelE | mRNA degradation | Dependent | A/T | (49) |
| BrnT | PF04365 | 15K | PF_BrnT | BrnT | mRNA degradation | Unknown | T/A | (80) |
| gp49 | PF05973 | 22K | PF_gp49 | gp49, gp49-like, tad, HigB1, HigB3, RelE, RelE/ParE | mRNA degradation | Dependent | T/A | (40, 48, 117) |
| DUF4258 | PF14076 | 6K | PF_DUF4258 | DUF4258, RelE | Unknown | Unknown | A/T | None |
| ParE-like | PF15781 | 2K | PF_ParE-like | RelE, ParE-like, RelE/ParE, RelE/StbE | Unknown but clustering analysis indicates mRNA degradation | Unknown | A/T | This work |
| ParE | PF05016 | 71K | PF_ParE | ParE, ParE1, ParE2, ParE3, ParE4, RelE, RelE1, RelE2, RelE3, RelG, RelK, StbE, YafQ, YoeB | mRNA degradation (RelE Cluster) & DNA Gyrase | Dependent (RelE Cluster) | A/T | (38, 61, 62, 113, 118) |

a) Protein names can be retrieved from InterPro using Pfam IDs;
b) Pfam holds in total 170K or ca. 170,000 BECR-fold toxins (unfiltered)
c) The list is not exhaustive;
d) GO, Gene Order; T = Toxin; A = Antitoxin;
e) Culviner, Laub et al (2021) found that MqsR cleavage of mRNA is effectively enhanced by the ribosome
e) Culviner, Laub et al (2021) found that YafO cleavage of mRNA might occur independently of the ribosome

| **Antitoxin - Toxin and Toxin - Antitoxin Gene Fusions Discovered** |  |  |  |  |
| --- | --- | --- | --- | --- |
| **Toxin Family** | **Antitoxin - Toxin Fusion** | **Prokaryotic** | **Eukaryotic** |  |
| **PF_YoeB** | YefM-YoeB fusions 7 (2) | 5 | 2 |  |
| **PF_YhaV** | PrlF-YhaV fusions 2 (1) | 1 | 1 |  |
| **PF_ParE** | ParD-ParE fusions 20 (1) | 19 | 1 |  |
|  | **Toxin - Antitoxin Fusions** |  |  |  |
| **PF_HigB** | HigB-HigA fusions 12 (2) | 10 | 2 |  |
| **PF_HigB-like** | HigB-HigA-like fusions 20 (3) | 17 | 3 |  |
| **PF_RelE** | RelE-YiaG fusions 7 | 7 | 0 |  |
| **PF_MqsR** | MqsR-MqsA fusion 5 | 5 | 0 |  |
| **PF_BrnT** | BrnT-BrnA fusions 32 (4) | 28 | 4 |  |
| **PF_gp49** | gp49-HTH fusions 69 | 69 | 0 |  |
| **PF_DUF4258** | DUF4258-MqsA fusions 17 | 17 | 0 |  |
| **PF_ParE-like** | ParE-like-Phd/YefM fusion | 1 | 0 |  |
| **PF_ParE** | ParE-ParD fusion | 3 | 0 |  |
| **Total** |  | 182 | 13 | 195 |

However, the classification and nomenclature of RelE/ParE superfamily toxins in the literature have often been inconsistent. For instance, toxins belonging to the RelE family (Pfam ID: PF06296) have been variously named HigB-2, RelE, RelE/ParE, or StbE, while the prototypical RelE of *E. coli* K- 12 is classified within the ParE family (Pfam ID: PF05016). To address this ambiguity, this work uses Pfam-based nomenclature for the 13 toxin families to improve clarity and consistency. For example, PF06296 is referred to as PF_RelE, and PF06769 as PF_YoeB (see **Table 1** for the complete list).

### The Pfam database does not separate RelE and ParE monodomain toxins into discrete families

The integration of Pfam into the InterPro Database (63) has enhanced accessibility to the 13 BECR-fold toxin families of the RelE/ParE superfamily. To analyze these families, individual Pfam protein families (PFs) were downloaded, and incomplete or multi-domain sequences were excluded. The curated dataset was reduced to approximately 9,000 representative sequences (**Table S1**) and subjected to clustering analysis.

In most cases, Pfam families clustered distinctly, with experimentally characterized toxins aligning to their respective clusters (**Figure 2**). For instance, all 10 experimentally validated ParE toxins localized within the PF_ParE cluster, and all four characterized YoeB toxins were found within the PF_YoeB cluster. Minor cross-cluster "spill-overs" were observed, such as a few PF_HigB sequences being found in the PF_YoeB cluster, but these occurrences were limited. Overall, the clustering largely agrees with Pfam annotations, with two notable exceptions.

**Figure 2.**
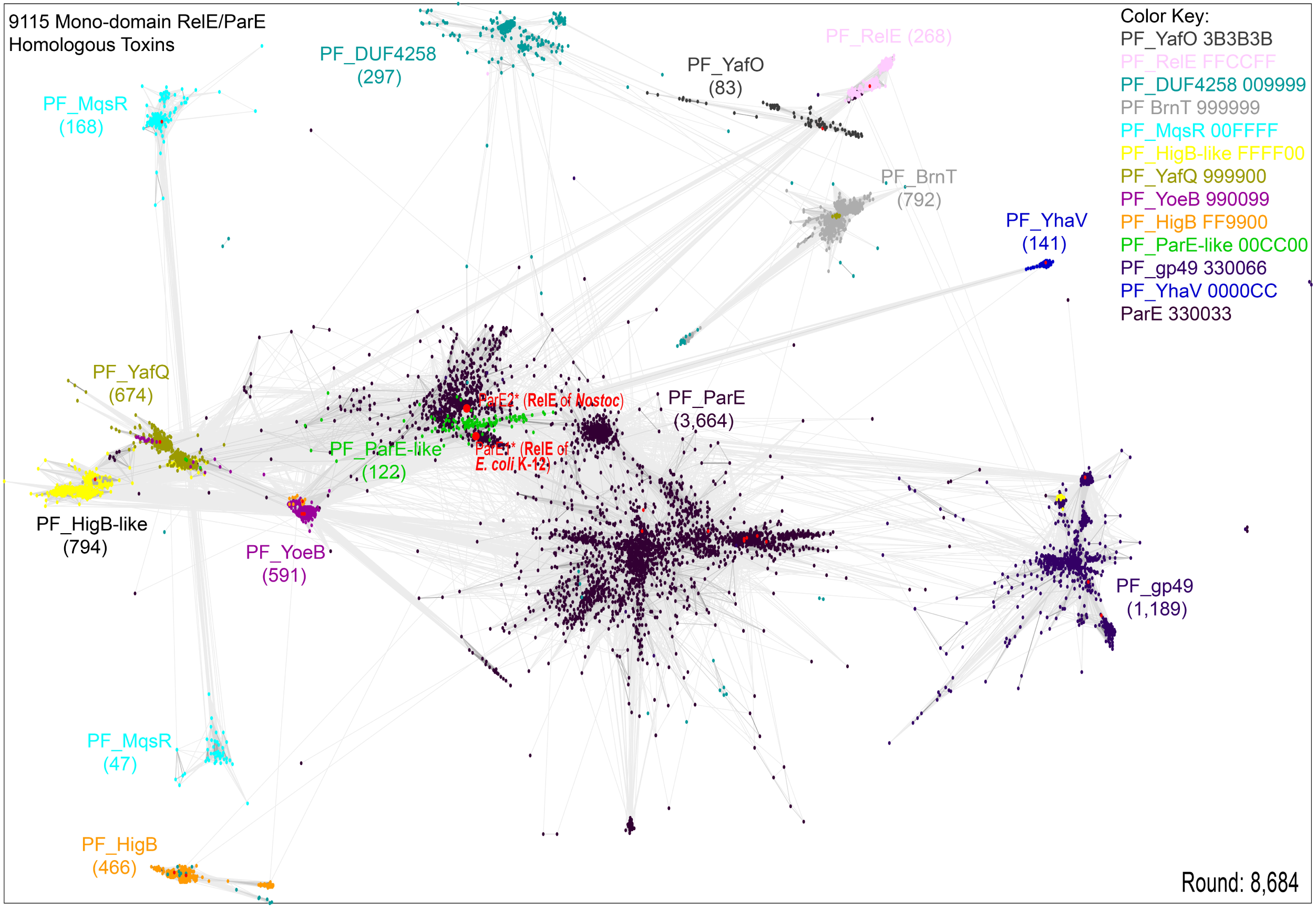
Clustering analysis of monodomain toxins from the RelE/ParE superfamily This 2D representation depicts a 3D clustering analysis of 9,115 toxin sequences from 13 Pfam families, all characterized by the BECR-fold. Each node corresponds to a single sequence, with connections between nodes representing significant pairwise sequence similarities. Number of toxins from each PF are given in parentheses; experimentally confirmed toxins are indicated with red dots. Note that the PF_ParE-like cluster group together with the subcluster of PF_ParE that contain the two ribosome-dependent RNases RelE of *E. coli* K-12 and Nostoc. Color-coding: Hex color codes: PF_YafO: 3B3B3B; PF_RelE: FFCCFF; PF_DUF4258: 009999; PF_BrnT: 999999; PF_MqsR 00FFFF; PF_HigB-like: FFFF00; PF_YafQ: 999900; PF_YoeB: 990099; PF_HigB: FF9900; PF_ParE-like: 00CC00; PF_gp49: 330066; PF_YhaV: 0000CC; PF_ParE: 330033.

The PF_ParE family, the largest BECR-fold toxin cluster, was unexpectedly found to split into several subclusters. Among these, one subcluster included ribosome-dependent RNases from *Escherichia coli* K-12 and *Nostoc*. These RNases differ functionally from other PF_ParE toxins, which inhibit DNA gyrase. This discrepancy highlights an inconsistency between Pfam classification and biological function (**Figure 2**). Notably, *E. coli* K-12 and *Nostoc* RelE toxins grouped within the same subcluster, reinforcing their shared function.

Another unexpected finding was the overlap of the PF_ParE-like cluster with the PF_ParE subcluster containing the ribosome-dependent mRNases. (Green cluster in **Figure 2**). This suggests that some PF_ParE-like toxins (PF15781) may also function as mRNases. Similarly, PF_MqsR toxins split into two clusters; however, this separation disappeared when a larger number of MqsR homologs were analyzed independently (data not shown).

### Phylogenetic analysis of PF_ParE and PF_ParE-like families

To address the observed incongruities, PF_ParE and PF_ParE-like families were analyzed together through clustering and phylogenetic approaches. PF_ParE was found to consist of at least five subclusters (**Figure S1** and **Table S1**). Phylogenetic analysis of these subclusters and PF_ParE-like toxins revealed distinct branching patterns with high statistical significance (**Figure 3A**). Consistent with the clustering results mentioned above (**Figure 2**), the PF_ParE-like toxins grouped within Subcluster #4, which contains the ribosome-dependent mRNases. A broader view of the phylogenetic tree (**Figure 3B**) shows PF_ParE-like and Subcluster #4 separating from the remaining four PF_ParE subclusters with high bootstrap support. This finding strengthens the hypothesis that both PF_ParE-like and Subcluster #4 toxins are mainly ribosome-dependent mRNases.

**Figure 3.**
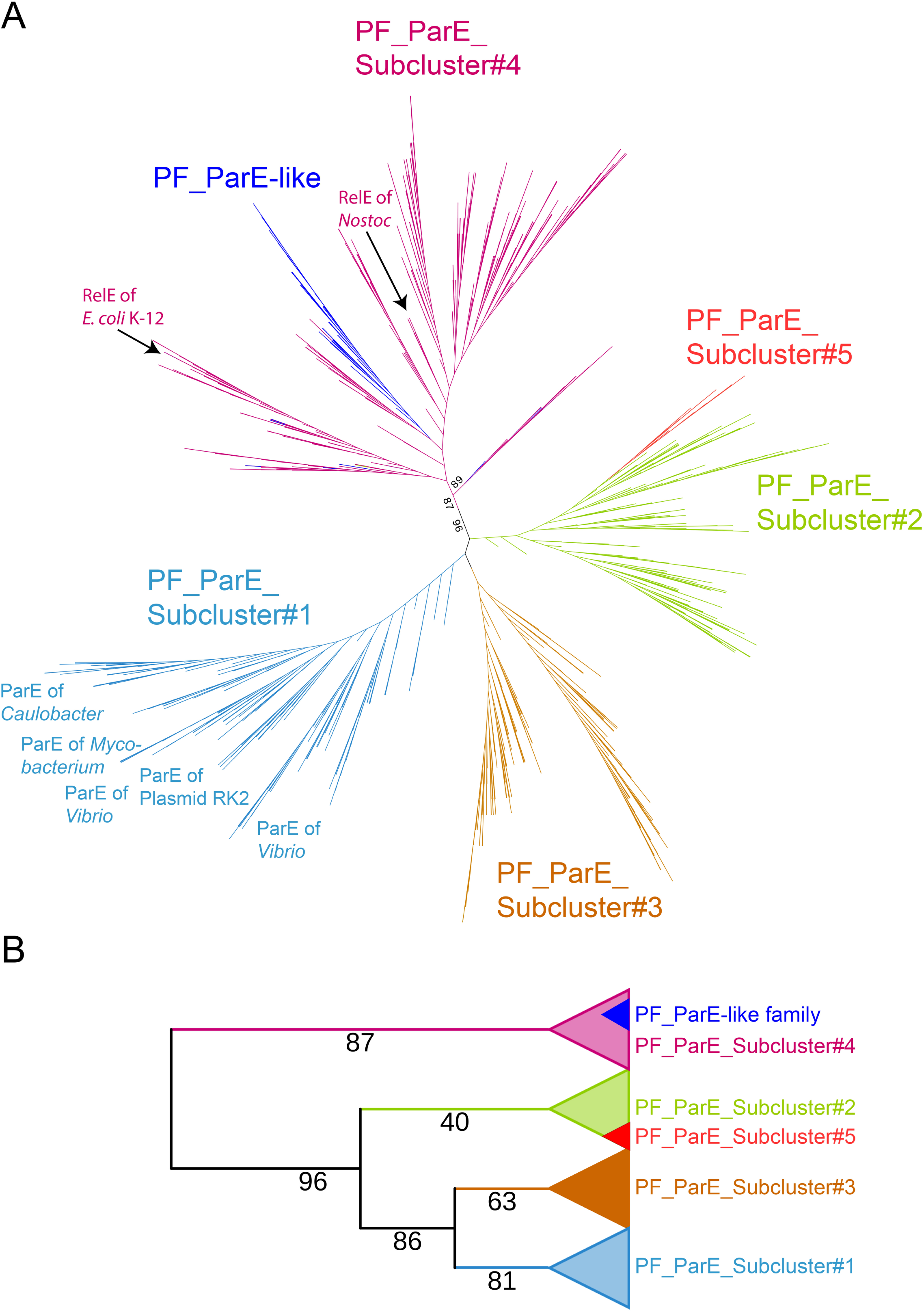
Phylogeny of the five subclusters of PF_ParE and PF_ParE-like toxins. A. Unrooted phylogenetic tree. The unrooted tree was constructed using FastTree from a ClustalO alignment of 1,913 sequences belonging to the PF_ParE and PF_ParE-like families. The tree highlights five distinct PF_ParE subclusters, as well as the PF_ParE-like family (blue). Notably, PF_ParE-like sequences are embedded within Subcluster #4, which also includes the ribosome-dependent mRNases RelE from *E. coli* K-12 and Nostoc. This suggests a functional and evolutionary link between the two groups. Additionally, Subcluster #1 predominantly comprises DNA gyrase inhibitors from various species, including *Caulobacter*, *Mycobacterium*, and plasmid RK2. B. Collapsed version of the tree. A simplified representation of the phylogenetic tree from (A) showing the relationships among the five subclusters and PF_ParE-like toxins. Bootstrap values are indicated on the branches, demonstrating high statistical support for the phylogenetic separation of subclusters. The bootstrap values highlight the robustness of clustering, with particularly high support for Subcluster #1 and the grouping of PF_ParE- like family within Subcluster #4.

### Note on DNA gyrase inhibitors in archaea

The analysis of DNA gyrase inhibitors (ParEs) across subclusters reveals a striking distribution bias, particularly in archaea. Subcluster #1 (432 toxins) and Subcluster #2 (378 toxins) contain no toxins originating from archaea, while Subcluster #3 (338 toxins) contains only a single archaeal toxin. In sharp contrast, Subcluster #4 includes 138 archaeal toxins out of a total of 734 toxins. The near-absence of DNA gyrase inhibitors in most archaeal subclusters aligns with the well-documented differences in their DNA topology management strategies. Unlike bacteria, which predominantly rely on DNA gyrase for introducing negative supercoils, archaea employ alternative mechanisms. Thermophilic archaea often utilize reverse gyrase, an enzyme unique to thermophiles that introduces positive supercoils to stabilize DNA at high temperatures. Conversely, mesophilic and psychrophilic archaea predominantly depend on topoisomerases. The clustering data reinforces the notion that bacterial-type DNA gyrase and its associated inhibitors, such as ParEs, are largely absent in archaea due to these alternative mechanisms. The presence of ribosome-dependent BECR- fold mRNases in archaea is well documented (39, 64, 65).

### Implications for functional annotation

The shared BECR-fold of RelE and ParE toxins (54) makes it challenging to predict whether a newly identified BECR-fold toxin functions as an mRNase or a DNA gyrase inhibitor. Phylogenetic approaches, such as those illustrated in **Figure 3A**, can provide valuable insights into the functional classification of uncharacterized BECR-fold toxins based on their position within the tree.

### Individual BECR-fold monodomain toxin families

The Pfam database holds thirteen BECR-fold toxin families encoded by operons potentially also encoding an adjacent antitoxin, amounting to approximately 170,000 potential toxins (**Table 1**). The number of toxins in each family is given in the table. It should be noted that these numbers include all kinds of potential BECR-fold toxins, including multidomain proteins, toxins encoded by solitary genes and incomplete toxins. After downloading, solitary and incomplete toxins were removed or, in the case of multidomain proteins, analyzed separately.

### PF_RelE

The PF_RelE family includes toxins often referred to as HigB in the literature, a naming convention that can lead to confusion. This likely stems from the inverted gene order of *higBA* (toxin upstream of antitoxin) compared to *relBE* (antitoxin upstream of toxin). Previously, we annotated and analyzed two toxin-antitoxin (TA) loci in the superintegron of *Vibrio cholerae*, both encoding RelE-homologous toxins (47). These loci were named *higBA-1* and *higBA-2* due to their gene order, which mirrors the *higBA* locus of the Rts1 plasmid, also encoding a RelE-homologous toxin (4, 66).

Similar to RelE of *E. coli* K-12, all three HigB toxins cleave mRNA codons at the ribosomal A-site (47). Likewise, these *higBA* operons are induced transcriptionally during amino acid starvation and, in addition, can stabilize plasmids (47, 66). Despite these similarities, the toxins belong to different Pfam families. Specifically, HigB-2 (*V. cholerae*, Q9KMA6) belongs to PF_RelE, while HigB-1 (*V. cholerae*, Q9KMG5) and HigB (Rts1, Q7A225) belong to PF_HigB-like [(63) Structurally, HigB-2 is more closely related to *E. coli* K-12 RelE than to HigB of Rts1, particularly with respect to the topology of the active site (67)]. The N-terminally disordered region of HigA-2 folds upon binding HigB-2, inducing a structural change of the active site and inhibiting the toxin’s catalytic activity (67).

### PF_YoeB

The *yefM-yoeB* TA module of *E. coli* K-12 encodes YoeB (P69348), an mRNA-cleaving toxin, and YefM, its antitoxin (68). Homologs such as Txe (Q848U6) from the *axe-txe* module of*nterococcus faecium* share similar properties. While YoeB cleaves RNA non-specifically in vitro without ribosomes (69), it cleaves mRNA codons at the ribosomal A-site in vivo, similar to RelE (40, 45, 46, 70). The YoeB/Txe family is widely distributed across bacterial phyla but is absent in archaea (**Table S1**).

### PF_HigB and PF_HigB-like

The *higBA* locus of the Rts1 plasmid, first identified for stabilizing plasmid inheritance, encodes HigB (Q7A225), a toxin that inhibits translation by cleaving mRNA at the ribosomal A-site (66, 71–73). Similarly, *E. coli* K-12 HigB (P64578) cleaves translated mRNAs, with its toxicity counteracted by HigA (24, 40, 47). We discovered *higBA* of *E. coli* K-12 and showed that HigB (P64578) cleaves translated mRNAs and that HigA counteracts HigB toxicity (57). Both HigA and the HigBA complex autoregulate the *higBA* through binding palindromic sequences in the promoter region (57, 74).

Three-component Toxin-Antitoxin-Chaperone (TAC) modules, which encode a SecB-like chaperone in addition to toxin and antitoxin, prevent degradation of their cognate antitoxins (75,76). HigB, encoded by the TAC module *higBAC* of *E. coli* C496, is activated by the λ phage major tail protein, thereby triggering abortive infection (24). The λ tail protein disrupts the interaction between HigA and the HigC chaperone, destabilizing HigA and activating HigB, thereby aborting phage replication.

### PF_MqsR

The *mqsRA* module, originally linked to *cspD* regulation, cleaves mRNA independently of ribosomes (57, 77). However, a recent study suggests that ribosomes enhance cleavage efficiency, challenging its absolute ribosome-independency (78). Some *mqsRA* modules are also part of TAC systems (79).

### PF_YafQ, PF_YhaV and PF_YafO

YafQ (Q47149) and YhaV (P64594) inhibit translation by cleaving mRNA at the ribosomal A-site, with YhaV favoring a G downstream of the cleavage site (49, 60). YafO (Q47157) may also exhibit ribosome-independent cleavage, but this requires further validation (78).

### PF_BrnT

BrnT (Q2YQ91), encoded by the *brnTA* locus of *B. abortus*, has in vitro ribonuclease activity but its in vivo dependence on ribosomes remains unclear (80).

### PF_gp49

Toxins in this family, often labeled as RelE or HigB, are ribosome-dependent mRNases (40, 48). Moreover, HigB of *E. coli* O112ab:H26 is part of a TAC module that functions in abortive infection defense (24).

### PF_DUF4258 and PF_ParE-like

Little is known about toxins in these families, as they remain undescribed in the literature.

### PF_ParE

PF_ParE is the largest BECR-fold family and includes both mRNases and DNA gyrase inhibitors. For example, RelE of *E. coli* K-12, a ribosome-dependent mRNase, paradoxically belongs to this family (**Figure 2**). Phylogenetic analysis of the subclusters derived from PF_ParE (**Figure S1**) suggests that Subcluster #1 primarily contains DNA gyrase inhibitors, while Subcluster #4 primarily includes mRNases (**Figure 3**).

### Multidomain proteins encoding BECR-fold toxin domains

To identify BECR-fold toxin domains in multidomain proteins, two complementary approaches were employed: (i) Sequence length analysis of the 13 Pfam toxin families include sequences significantly longer than the average monodomain length (**Table S2**). These extended sequences were filtered for further analysis using structural and sequence-based tools, including AlphaFold2, Foldseek, Phyre2, BLAST, and Hmmsearch (see *Materials & Methods*). This strategy revealed candidate multidomain proteins containing BECR-fold toxin domains embedded within larger architectures. (ii) Query-based searches for multidomain proteins. Monodomain toxins were used as queries in BLAST and Hmmsearch analyses to screen for related sequences in protein databases.

These searches successfully identified additional multidomain proteins containing BECR-fold toxin domains (**Table S2**).

It should be noted that many multidomain BECR-fold proteins exhibiting sequence similarity to members of the RelE/ParE family were exempted from the analysis presented here, either because an antitoxin gene or domain could not be identified or because there were too few members in a given multidomain protein family to yield a meaningful bioinformatics analysis.

### Fused toxin – antitoxin mono-gene modules

Several examples from the literature show that TA gene fusions can encode functional protein fusions and that some of them function in phage defense (42, 81). In total 195 genes encoding a BECR-fold domain fused to an antitoxin domain were identified (**Table 2; Table S2).** That these genes are so common indicate that at least some of them are functional. Below are comments on the individual fused TA families identified here. It should be noted that the CARF-RelE toxin family and its activation mechanism was recently described elsewhere (42).

### Mono and bicistronic *relE yiaG* modules

YiaG is a Helix-Turn-Helix (HTH) transcription factor of *E. coli* K-12 with a C-terminal XRE-type HTH DNA-binding domain. Transcription of *yiaG* is upregulated in response to stress, the stringent response, and glucose-lactose diauxic carbon-source shifts (82). However, its function is unknown.

Data in **Table S1** reveal that a subfamily of PF_RelE toxins with extended N-termini are encoded by genes positioned upstream of *yiaG* homologs (**Figure 4A**). Sequence alignments of these extended N-terminal regions show high heterogeneity (**Figure 4B**) and structural modeling predicts that these regions are intrinsically disordered. As an example, modeling of RelE of *E. coli* VREC0535 reveals its C-terminal BECR-fold domain and its N-terminal unstructured domain (**Figure S2A**).

**Figure 4.**
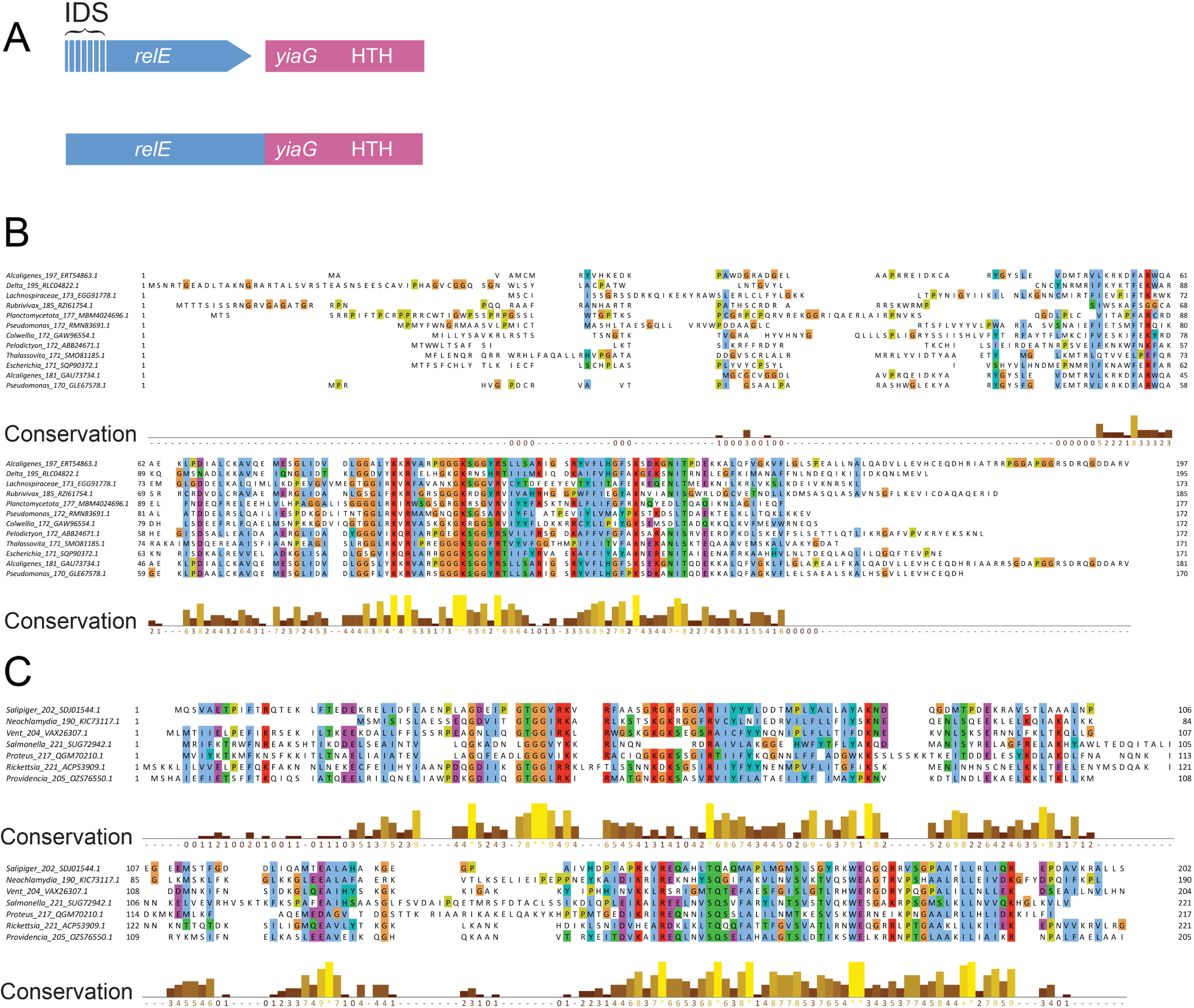
Features of mono and bicistronic *relE yiaG* toxin-antitoxin modules. A. Genetic organization of mono (fused) and bicistronic *relE yiaG* modules. The bicistronic module encodes two proteins: RelE (toxin) and YiaG (antitoxin), of which the latter containing a C-terminal HTH domain. In monocistronic modules, the *relE* and *yiaG* genes are fused into a single protein with a RelE-like toxin domain and an antitoxin domain homologous to YiaG. The hatched region in RelE denotes the unstructured or intrinsically disordered segment. B. Sequence alignment of PF_RelE toxins with extended N-termini. The alignment highlights the conserved regions and variability of the N-terminal intrinsically disordered segments across PF_RelE toxins associated with YiaG antitoxins. C. Sequence alignment of RelE-YiaG fusion proteins, highlighting conserved residues and structural motifs across diverse organisms. Conservation scores are indicated below the alignment.

Modeling of YiaG from *E. coli* VREC0535 reveals its C-terminal HTH domains connected to two parallel helices connected by an unstructured linker (**Figure S2B, upper panel**). Database search via revealed that the HTH domain of YiaG closely resembles that of the antitoxin HigA2 of *V. cholerae* (**Figure S2B, lower panel**). Furthermore, structure modeling predicts that two RelE•YiaG heterodimers dimerize via YiaG’s HTH domains (**Figure S2C**). The interaction between YiaG’s N- terminal α-helices and the C-terminal α-helices of the RelE BECR-fold domain suggests a possible inhibitory mechanism through this interface.

### Fused *relE-yiaG* operons

Monocistronic operons encoding RelE-YiaG fusion proteins were also identified (**Table S2**). Alignments of RelE-YiaG fusions reveal two conserved domains linked by non-conserved regions (**Figure 4C)**. Modeling of the RelE-YiaG fusion protein from *Salmonella enterica* does not indicate direct interactions between the N-terminal RelE domain and the C-terminal YiaG domain in monomeric form (**Figure S2D**). However, structural overlays show striking similarity between the antitoxin domain of the fusion protein and YiaG from *E. coli* VREC0535, including similar α- helices in the N-terminus of YiaG and the N-terminal part of the antitoxin domain of the RelE-YiaG fusion (**Figure S2E**).

### Fused yefM-yoeB and RHH-yoeB operons

A YoeB toxin domain can be fused to an N-terminal YefM domain in a YefM-YoeB configuration (**Figure S3A; Table S2**). Phd/YefM (hereafter YefM) DNA-binding domains adopt a β1-α1-α2-β2-β3-α3 topology (83), which is preserved in the fusion protein from *Mycobacterium shinjukuense* (**Figure S3B**). Structural predictions suggest that the YefM-YoeB fusion dimerizes via its YefM domain, akin to other DNA-binding transcriptional regulators (**Figure S3C**). These proteins are predicted to bind operator sequences within the promoter region, consistent with their presumed transcriptional regulatory role. Overlaying the experimentally determined YefM structure from *E. coli* K-12 (69) with the predicted YefM structure from *M. shinjukuense* confirms the validity of the model (**Figure S3D**). In other cases, the YoeB domain is fused to a RHH domain, forming a RHH- YoeB configuration (**Table 2, Figure S3E**).

### Fused *BrnTA* modules in pro and eukaryotes

The two-gene operon *brnTA* of *Brucella abortus* encodes the BECR-fold toxin BrnT and its antitoxin BrnA (80). BrnA contains a C-terminal RHH domain and controls transcription of *brnTA* (**Figure 5A)**. This study uncovered *brnTA* modules encoding fusion proteins between BrnT and BrnA, with 31 out of 33 fusions containing RHH domains and 2 containing HTH domains (**Table 2, Table S2, Figure 5B**). Surprisingly, four of the TA fusions were from eukaryotic organisms, including fungi (Gigaspora), porifera (seasponges), and nematodes.

**Figure 5.**
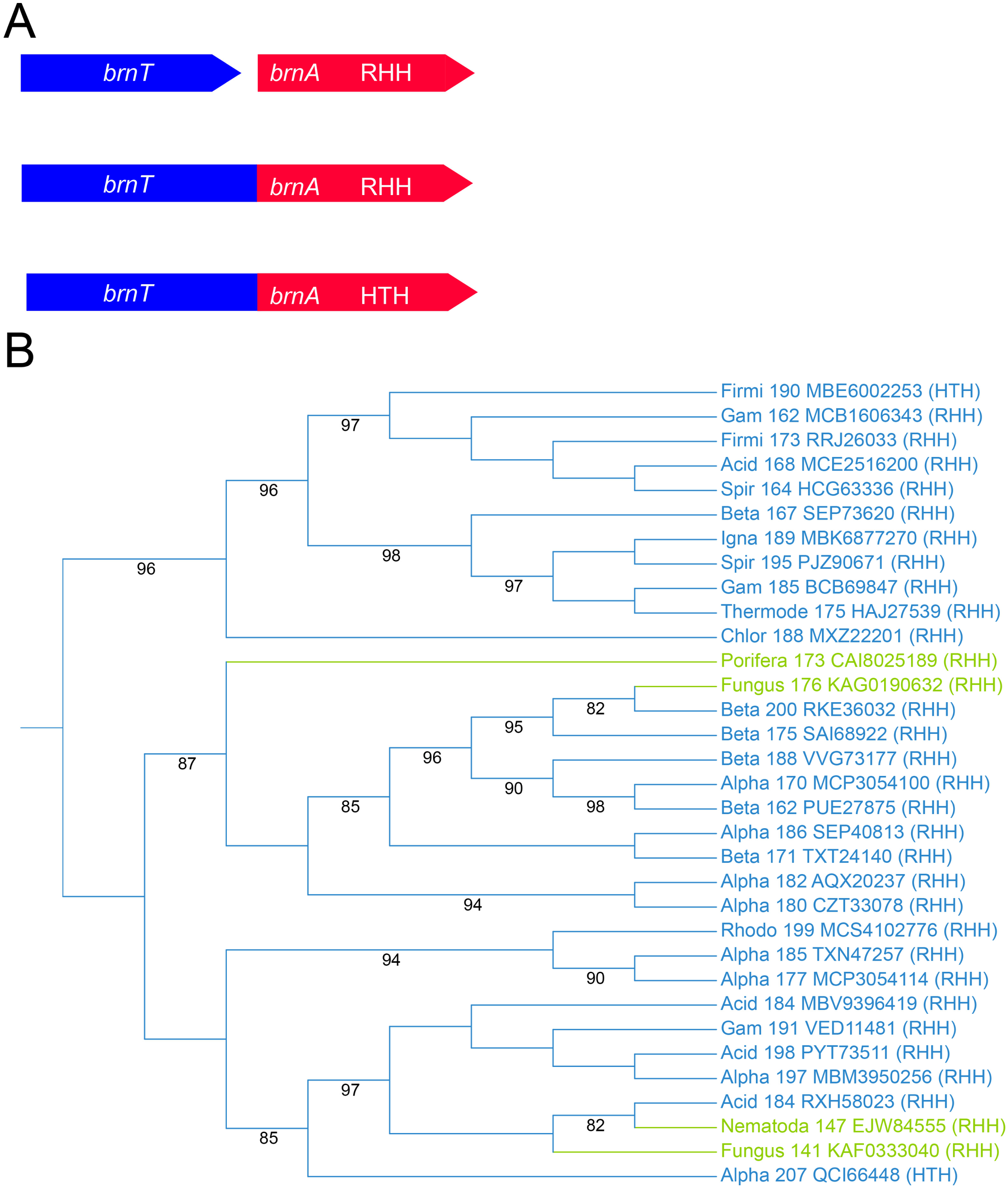
*BrnTA* fusions are present in both bacteria and eukaryotes A. Genetic organization of canonical bicistronic *brnTA* modules encoding an antitoxin with a RHH domain (top), *brnTA* fusion modules encoding an RHH domain or an HTH domain antitoxin (middle, bottom). B. Phylogenetic tree of *BrnTA* fusions from bacteria (blue) and eukaryotic organisms (green). Type of DNA-binding domain in the antitoxin part of the proteins is indicated (RHH or HTH). Bootstrap values > 80 are shown.

The presence of *brnTA* fusions in eukaryotes raises the possibility of sequence contamination in databases. However, it more likely reflects genuine genetic presence, either within the host genome or via symbiotic microbiota or endosymbionts. Phylogenetic analysis supports this notion, as eukaryotic *brnTA* fusions cluster non-randomly in the tree (**Figure 5B**). Additionally, these organisms are known to harbor endobacterial populations that facilitate horizontal gene transfer. This highlights the potential evolutionary and functional significance of *brnTA* fusions in diverse domains of life (84–87).

### Novel potential defence locus *ptuXYZ* encoding a BECR-fold RNase

The *ptuXYZ* operon encodes a conserved and widespread three-gene module with putative roles in anti-phage defense (**Figure 6**; **Table S2**). The third gene, *ptuZ*, encodes a BECR-fold RNase related to YoeB. Each component of the operon exhibits features characteristic of defense systems targeting invading foreign genetic material: The first gene, *ptuX*, encodes a protein with a DUF262 domain that is also found in Type IV restriction endonucleases such as GmrSD, SspE, and BrxU. These enzymes degrade epigenetically modified phage or plasmid DNA (88–90). Unlike Type IV restriction nucleases, which also contain a DUF1524 HNH nuclease domain, PtuX lacks this C- terminal domain, resulting in a shorter protein (**Table S2**). Structural comparisons reveal that PtuX resembles ParB/Srx nucleases (90, 91), consistent with its putative nuclease function. It should be noted that PtuX does not resemble PtuB.

**Figure 6.**
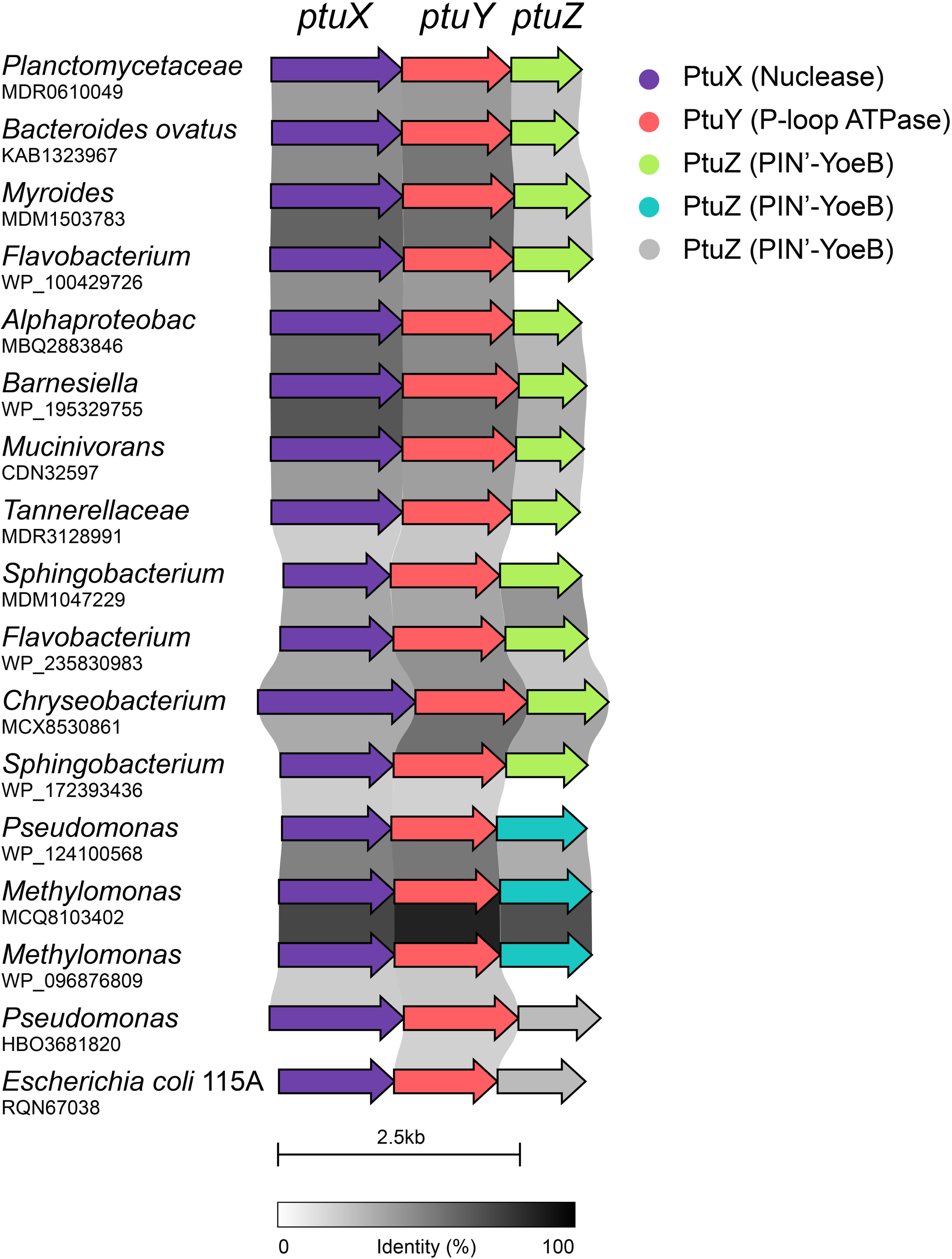
Schematic of a conserved defence operon encoding a BECR-fold RNase. The Figure illustrates 17 homologous operons, here collectively referred to as *ptuXYZ*. Each operon encodes three genes with predicted functions in anti-phage defence: PtuX contains DUF262, a domain shared with type IV restriction endonucleases such as GmrSD, BrxU, and SspE, which are involved in degrading epigenetically modified phage DNA (90, 91). PtuY exhibits structural similarity to PtuA, a P-loop ATPase serving as the regulatory component in the *ptuAB* septu anti-phage system (94). PtuZ is a two-domain protein comprising an N-terminal PIN-like domain and a C-terminal BECR-fold domain related to YoeB toxins. Genera and corresponding GenBank IDs of the operon’s first gene are listed for each organism. Operon relationships are visualized by homology shading between adjacent genes, with the gradient reflecting sequence identity. The schematic was generated using clinker (103) and is drawn to scale. Details of the protein sequences and database identifiers are provided in **Table S2**. Color coding of arrows: PtuX (Nuclease): Purple; PtuY (P-loop ATPase): Red; PtuZ (PIN’-YoeB variants): Light Green, Cyan and Pink. Scale bar: 2.5 kb. Sequence identity is indicated by the grayscale gradient (0–100%).

The second gene, *ptuY*, encodes a protein with an N-terminal P-loop ATPase domain. Structural alignments reveal significant similarity between the ATPase domains of PtuY and PtuA, with an RMSD of approximately 1 Å over 100 residues (**Figure S4**). PtuA is a component of Septu antiphage systems (92, 93), and the structural similarity thus reinforces that possibility that PtuY also functions in anti-phage defence.

Like PtuA, which forms a horseshoe-shaped trimer of dimers (94), structural modeling predicts that PtuY forms dimers (**Figure S5A**, **B**) and higher-order trimers-dimer complexes (**Figure S5C-F)**.

These closed-ring assemblies are reminiscent of prokaryotic AAA+ ATPases, such as ClpX, ClpA, and Lon proteases, which function in protein remodeling and degradation (95, 96).

The third gene, *ptuZ*, encodes a two-domain protein connected by a helical linker (**Figure 7A**). The C-terminal domain adopts a BECR-fold similar to that of HigB/YoeB toxins (**Figure 7C**), while the N-terminal domain resembles a VapC PIN-like RNase fold (**Figure 7B**). The PtuZ PIN-like domain lacks essential catalytic residues, suggesting it may serve a structural rather than an enzymatic role. The combination of nuclease (PtuX), ATPase (PtuY), and a BECR-fold RNase-like domain (PtuZ) strongly suggests that the *ptuXYZ* operon functions as a coordinated defense mechanism against foreign genetic elements, such as phage or plasmid DNA. Further experimental validation is needed to elucidate its precise molecular mechanism.

**Figure 7.**
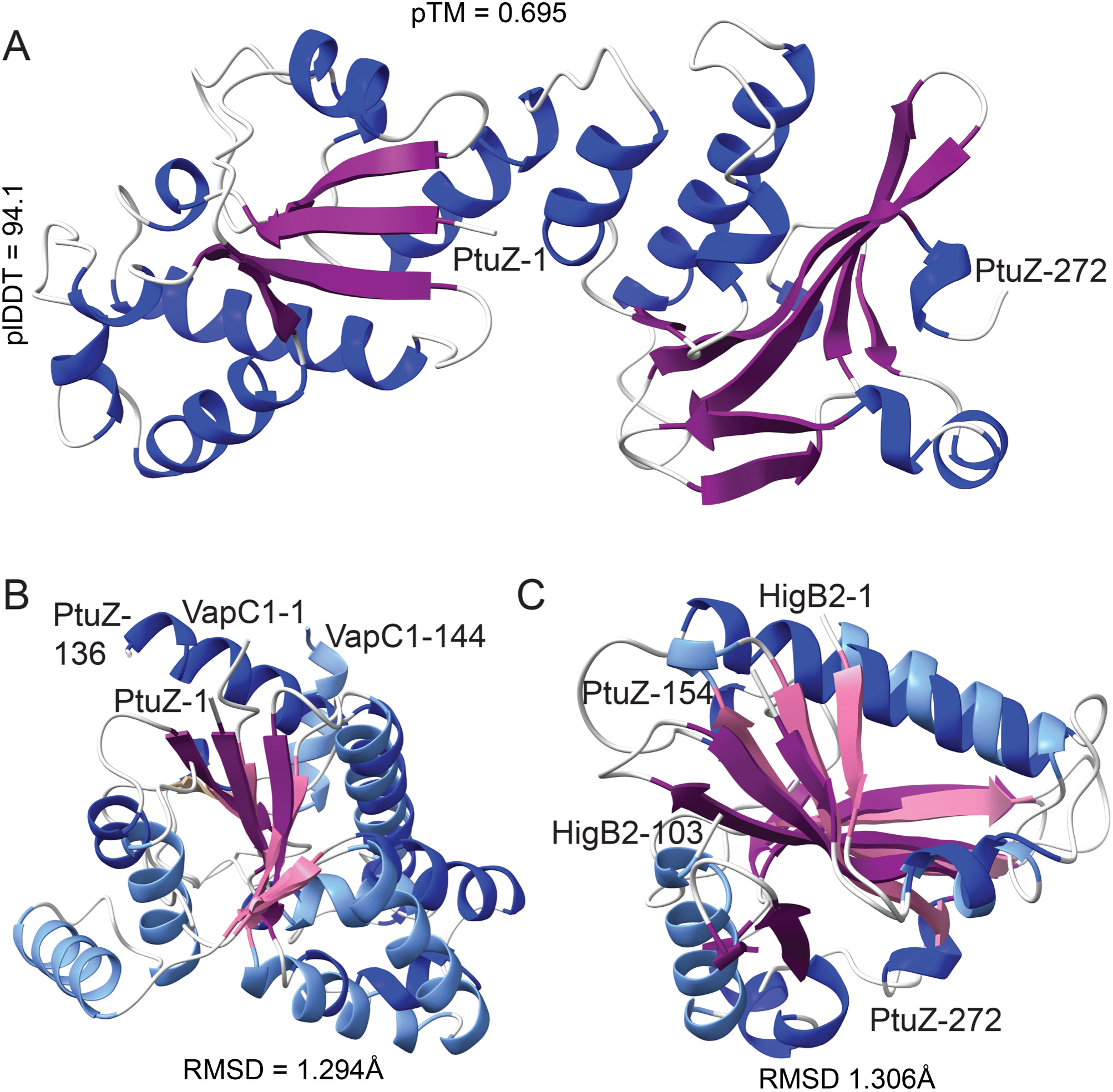
Structure of PtuZ and comparison with PIN and BECR-fold domain RNases. **A.** AF2-generated structural model of PtuZ from *Flavobacterium* sp. 1 (WP_100429724.1). The model quality metrics include plDDT = 94.1 and pTM = 0.695. Helices are represented in blue, while beta-sheets are colored purple. N- and C-terminal residues are labeled to indicate the protein’s domain boundaries. **B.** Superimposition of the N-terminal PIN-like domain of PtuZ and the PIN domain of VapC1 from *Aquifex aeolicus* VF5 (O66399.1). The structural alignment shows high similarity, with an RMSD of 1.294 Å over 25 pruned atom pairs. PtuZ domains are colored as in (A), while VapC1 domains are represented with light blue helices and pink beta-sheets. **C.** Superimposition of the C-terminal BECR-fold domain of PtuZ and HigB2 toxin from *E. coli* APEC179 (WP_032269392.1). The structural alignment yields an RMSD of 1.306 Å over 27 pruned atom pairs. PtuZ helices and sheets are colored blue and purple, respectively, while HigB2 is represented in light blue and pink.

### ToxM: A BECR-fold toxin with a transmembrane domain

BECR-fold toxins fused to transmembrane (TM) domains represent a unique configuration of type II TA systems. These toxins, here referred to as ToxM (for membrane toxin), exhibit a tripartite structure consisting of: (i) an intracellular N-terminal BECR-fold domain; (ii) a central TM domain predicted to span the membrane; (iii) an extracellular disordered region capped with well-structured C-terminal domain. This structural organization is exemplified by ToxM from *Xanthomonas perforans* (**Figure 6A**, **6B**). AF2 and Membranefold modeling predicts that the N-terminal BECR- fold domain is cytoplasmic, that the TM region very likely is embedded in the inner membrane and that the C-terminal domain is periplasmic (**Figures S6C**).

### ToxR: Antitoxins associated with ToxM

The *toxM* genes are co-transcribed with upstream *toxR* genes that encode small antitoxins containing RHH or HTH DNA-binding domains. These antitoxins likely regulate the activity of ToxM and transcription of *toxRM*. AF2 modeling predicts that ToxR of *X. perforans*, an RHH- domain regulator, forms dimers, consistent with the oligomerization properties of RHH domains (**Figure S6D**). These dimers are likely involved in DNA binding for transcriptional autoregulation, as commonly observed for Type II antitoxins. Modeling of ToxM_2_-ToxR_2_ tetramers suggests that ToxR antitoxins wrap around the N-terminal BECR-fold domains of their cognate ToxM toxins (**Figure S6E**). Interestingly, the ToxM domains within the complex do not interact directly, implying that the inhibitory mechanism of ToxR primarily involves physical occlusion or conformational constraints on ToxM catalytic activity. The functional roles of the ToxM-ToxR modules remain speculative but may involve interactions with extracellular or membrane-associated targets. It is thus possible that ToxMR is activated by passage of foreign DNA (e.g. phage or plasmid) through the inner membrane and thereby function to neutralize the intruder. To my knowledge, this is a first possible example of a membrane-associated type II TA complex.

## Conclusion

This study provides a comprehensive bioinformatics and structural analysis of BECR-fold toxins within Type II toxin-antitoxin (TA) systems, shedding light on their diversity, structural features, and potential functional roles. By systematically analyzing 13 distinct monodomain BECR-fold toxin families and extending the scope to multidomain and transmembrane-associated variants, this work uncovers important evolutionary and functional insights into the RelE/ParE superfamily.

The identification of novel multidomain BECR-fold toxins and the clustering of these proteins into phylogenetically distinct subgroups reinforce the idea that sequence and structural variation are tightly linked to functional specialization. The discovery of *ptuXYZ* operons and the structural modeling of their encoded components highlight their potential involvement in defense mechanisms against invading genetic elements, particularly phages or plasmids. Furthermore, the ToxM-ToxR modules reveal an intriguing new dimension to TA systems, wherein membrane localization and complex antitoxin-toxin interactions may contribute to defence against foreign DNA intrusion or other biological processes.

Importantly, the incongruities in the functional annotation of BECR-fold toxins, particularly in the overlapping RelE/ParE classifications, underscore the need for more robust experimental characterization to validate and refine these findings. The phylogenetic and clustering approaches developed in this study offer a practical framework for delineating toxin function and classification, opening the way for the systematic exploration of BECR-fold TA systems in diverse bacterial species. Together, these findings advance our understanding of BECR-fold toxins as critical players in bacterial defense and physiology. Experimental validation of these predictions, combined with exploration of their ecological and physiological roles, will be pivotal in unraveling the broader implications of TA systems in bacterial biology.

## Materials and Methods

### Clustering analysis of BECR-fold toxin families

To analyze these families, individual Pfam protein families (PFs) were downloaded, and incomplete or multi-domain sequences were excluded. The curated dataset was reduced to approximately 9,000 representative sequences (**Table S1**) and subjected to clustering analysis using CLANS (97). Sequences of the 13 BECR-fold toxins were downloaded from InterPro Database and reduced proportionally to ca. 9,000 sequences. These were then used in the CLANS analyses (**Figure 2, Table S1** and **Figure S1**).

### Sequence alignments and phylogenetic trees

Sequence alignments were generated by Clustal Omega (98) at www.ebi.ac.uk and imported into Jalview (99). Protein sequence alignments in Jalview 2.11.0 were exported as vector files (EPS or SVG formats), converted to jpg files and imported into Adobe Illustrator, annotated and saved in PDF format for publication. Phylogenetic trees were visualized using iTOL (100). Reconstruction of phylogenetic trees was accomplished using FastTree (101) via the CIPRES module in Genious Prime that uses the Maximum Likelihood

approach and Ultrafast bootstrapping.

### Toxin – antitoxin gene organization, gene neighborhood analysis and gene cluster analysis

were accomplished using webFlaGs (102), TADB3.0 (55) and clinker (103). The gene organizations shown in the Figures were then confirmed by manual inspection of the DNA regions encoding BECR-fold toxins of **Table S1**.

### Protein Structure prediction, protein similarity searches and, protein structure visualization

Protein tertiary structures were modelled using AlphaFold2 (104) via the ColabFold v1.5.2-patch (105) or the AlphaFold patch of ChimeraX. Transmembrane toxins were analysed using Membranefold (Jeppe Hallgren, Ole Winther et al., bioRxiv, 2022). Mutimeric structures were modelled by MultiFOLD (106) or AlphaFold2 and validated by ModFOLDdock (107). Structure similarity searches was done using Phyre2 (108) or Foldseek (109). Structures were visualized and annotated using ChimeraX (110). HTH motifs were identified by EMBOSS (111).

